# Sugar metabolism of the first thermophilic planctomycete Thermogutta terrifontis: comparative genomic and transcriptomic approaches

**DOI:** 10.1101/179903

**Authors:** A.G. Elcheninov, P. Menzel, S.R. Gudbergsdottir, A. Slesarev, V. Kadnikov, A. Krogh, E.A. Bonch-Osmolovskaya, X. Peng, I.V. Kublanov

## Abstract

Xanthan gum, a complex polysaccharide comprising glucose, mannose and glucuronic acid residues, is involved in numerous biotechnological applications in cosmetics, agriculture, pharmaceuticals, food and petroleum industries. Additionally, its oligosaccharides were shown to possess antimicrobial, antioxidant and few other properties. Yet, despite its extensive usage, little is known about xanthan gum degradation pathways and mechanisms.

*Thermogutta terrifontis* R1 was described as the first thermophilic representative of the *Planctomycetes* phylum. As other cultivated planctomycetes, it grows well on various carbohydrates including oligo- and polysaccharides, however, its capability of anaerobic growth with or without electron acceptors was a novel finding among the representatives of this phylum.

The aim of this work is to examine *T. terrifontis* catabolic pathways with a special focus on the xanthan gum degradation pathway using genomic and transriptomic sequencing. Genomic analysis revealed more than a hundred glycosidases, polysaccharide lyases and other CAZymes, involved in oligo- and polysaccharide degradation by *T. terrifontis*, proteins of central carbohydrate metabolism and aerobic and anaerobic respiration. Furthermore, the combination of genomic and transcriptomic approaches revealed a putative novel xanthan gum degradation pathway involving unusual catalytic steps and enzymes: novel glycosidase(s) of DUF1080 family, hydrolyzing xanthan gum beta-glucosidic backbone linkages and beta-mannosidases instead of xanthan lyases for degradation of terminal beta-mannosidic linkages. Surprisingly, the genes coding DUF1080 proteins were found in high number in *T. terrifontis* and in many other *Planctomycetes* genomes, which, together with our observation that xanthan gum being a selective substrate for many planctomycetes, supports the important role of DUF1080 in xanthan gum degradation. Our findings shed light on the metabolism of the first thermophilic planctomycete, capable to degrade a number of polysaccharides, either aerobically or anaerobically, including the biotechnologically important bacterial polysaccharide xanthan gum. The results serve as good foundation for future exploration of *T. terrifontis* and its enzymes in biotechnological applications.

## 1. Introduction

Although 78,431 16S ribosomal RNA sequences are assigned to bacterial phylum *Planctomycetes* in the SILVA Ref database (release 128, Quast et al., 2013), only few tens cultivated species are described, and 38 genome sequences of cultivated and ca. twice that amount of uncultivated planctomycetes are available in the IMG database (March 2017, Markowitz et al., 2016). Altogether, the cultivated planctomycetes with validly published names comprise two classes, three orders, five families, 25 genera and 29 species. Until recently, all of these were characterized as strictly aerobic, heterotrophic and peptidoglycan-less microorganisms, which reproduce by budding and grow at mesophilic and slightly psychrophilic conditions (Fuerst and Sagulenko, 2011; Liesack et al., 1986). However, all of these features were reconsidered during the past few years. Their cell walls have been shown to contain a uniquely thin peptidoglycan layer (Jeske et al., 2015), representatives of the novel class *Phycisphaerae* divide by binary fission (Fukunaga et al., 2009; Kovaleva et al., 2015) instead of budding, and, finally, a few thermophilic and facultative anaerobic representatives were recently isolated (Slobodkina et al., 2015; Kovaleva et al., 2015). Even though no autotrophic planctomycetes were isolated and cultivated so far, members of the third class-level lineage, represented by uncultivated anammox planctomycetes (van de Graaf et al., 1995), are thought to fix CO2 via the acetyl-CoA pathway (Strous et al., 2006).

*Thermogutta terrifontis* R1 was characterized as the first thermophilic representative of the phylum *Planctomycetes* (Slobodkina et al, 2015). Moreover, it was among the first anaerobically grown planctomycetes and the first one growing by anaerobic respiration. However, nothing is known on the mechanisms underlying these novel capabilities. *Thermogutta terrifontis* strain R1 has been shown to grow on xanthan gum (Slobodkina et al, 2015) – a complex polysaccharide synthesized by *Xanthmonas campestris*, comprising a beta-1,4-glucan backbone and mannosyl–glucuronyl–mannose side chains. Xanthan gum itself has numerous biotechnological applications in cosmetics, agriculture pharmaceuticals, food and petroleum industries (Garcia-Ochoa et al., 2000). Moreover, its oligosaccharides were described as elicitors, stimulating plants in their defense response against pathogens (Liu et al., 2005); as antimicrobial compounds (Qian et al., 2006) and as antioxidants (Xiong et al., 2013). At the moment, not much is known about its decomposition pathways, especially on mechanisms and involved proteins. A few key enzymes needed to break xanthan gum side chains (xanthan lyases, beta-glucuronyl hydrolases and alpha-mannosidases) are known, yet still no glycosidases acting on the glucan backbone of xanthan gum have been characterized. Here, we reconstruct the central carbohydrate metabolism of *Thermogutta terrifontis* strain R1 using genomic and transcriptomic sequencing with a special emphasis on xanthan gum degradation.

## 2. Materials and Methods

### 2.1. Cultures and DNA / RNA extraction

*T. terrifontis* R1 cells were grown under microaerophilic conditions for 2-4 days in glass bottles, sealed with butyl rubber plug with aluminium cap, on modified Pfenning medium supplemented with xanthan gum or trehalose as a growth substrate. Mineral growth medium (Podosokorskaya et al., 2011) was prepared aerobically; 50 mg l^-1^ of yeast extract (Sigma) was added as a source of indefinite growth factors. Atmospheric air was in the gas phase and no reducing agents or resazurin were added. pH was adjusted to 6.1 with 5 M NaOH. Xanthan gum (1 g/l) (KELTROL^®^T, (food grade Xanthan Gum, Lot#2F5898K) CP Kelco) or trehalose (2 g/l) (Sigma, T9531) were added from sterile stock solutions before inoculation.

For genome sequencing 2000 ml of *T. terrifontis* grown culture was centrifuged at 10000 rpm, and cells pellet was harvested. Total DNA was extracted from the cell pellet by freezing and thawing in TNE buffer (Tris 20 mM, NaCl 15 mM, EDTA 20 mM). After treating with lysozyme, RNase A, SDS and proteinase K the DNA was extracted with phenol/chloroform and precipitated with EtOH and dissolved in 2 mM TE buffer (Gavrilov et al., 2016).

For the transcriptomic experiment, cultures on both xanthan gum and trehalose were grown for 4 days. Then, three samples of each culture (cultivated on xanthan gum and trehalose) were used for RNA extraction. RNA was extracted using TRI-reagent (Sigma), following standard protocol with the addition of two freeze and thaw cycles of the cells in TRI-reagent, as well as addition of chloroform and washing twice with 75% EtOH. RNA pellet was dissolved in RNAase-free water.

### 2.2. Sequencing, Assembly, and Mapping

The genome was sequenced using a combination of Illumina GA II, and Roche/454 sequencing platforms. All general aspects of 500 bp paired-end (PE) library construction and sequencing, as well as 454 single-end sequencing can be found at the corresponding company website. A hybrid assembly of Illumina and 454 datasets was done using the Newbler assembler (Margulies et al., 2005). The initial Newbler assembly consisting of 7 unique contigs was used to identify repeat regions that were subsequently screened out. At this point a 2 kb mate-pair Illumina library was constructed and sequenced and obtained paired end information was used to arrange multiple screened contigs into a single scaffold using the Phred/Phrap/Consed software package (Gordon et al., 1998). This package was also used for further sequence assembly and quality assessment in the subsequent finishing process. Sequence gaps between contigs that represented repeats were filled with Dupfinisher (Han and Chain, 2006), and a single scaffold was manually created and verified using available paired-end information. Illumina reads were used to correct potential base errors and increase consensus quality. Together, the combination of the Illumina and 454 sequencing platforms provided 320× coverage of the genome.

### 2.3. Genome Annotation

The assembled chromosome was uploaded to the RAST server (Aziz et al., 2008) for *de novo* gene prediction using Glimmer-3 (Delcher et al., 2007)) and initial detection of homologs. Furthermore, the predicted genes were searched against the protein databases Pfam 27.0 (Finn et al., 2014), COG 2003-2014 (Galperin et al., 2015), MEROPS 9.12 (Rawlings et al., 2016), and TCDB (Saier et al., 2014) in order to expand the initial RAST annotation. Signal peptides were predicted with the SignalP 4.1 web server (Petersen et al., 2011), and transmembrane helices were predicted with the TMHMM 2.0 web server (Krogh et al., 2001). Infernal 1.1.1 (Nawrocki and Eddy, 2013) was used in conjunction with the covariance models from Rfam 12.0 (Nawrocki et al., 2015) to search for non-coding RNA genes.

### 2.4. Transcriptome sequencing and assignment

Extracted total RNA was converted to cDNA by reverse transcriptase. Total cDNA was sequenced from the 3 samples of each culture by strand-specific paired-end Illumina sequencing using an insert size of 270 bp and read length of 90 bp. RNA-seq reads were mapped to the genome using BWA ver 0.7.8 (Li and Durbin, 2009) requiring properly mapped pairs. Read assignments to genes was accomplished using the featureCounts program from the Subread package (Liao et al., 2013). Only uniquely assigned read pairs were counted. Differential gene expression between the two groups was measured using edgeRun (Dimont et al., 2015) and genes were called differentially expressed using a BH-corrected p-value of 0.05.

### 2.5. Plylogenetic analysis.

Phylogenetic analyses were performed according to Sorokin et al., 2016 using the maximum likelihood method in MEGA6 (Tamura *et al*., 2013). Initial multiple amino acid sequence alignments were done in Mafft 7 (Katoh et al., 2013).

## 3. Results and discussion

### 3.1. Genome assembly and general genome characteristics

The genome of *Thermogutta terrifontis* strain R1 was sequenced and assembled into a single circular chromosome with a length of 4,810,751 bp and GC content of 57.34%. Genome annotation was performed using the RAST server and Infernal. In total, 4,504 protein coding genes were found in the genome, of which 2,412 could not be annotated by our database search and are therefore designated as “hypothetical protein”.

Both RAST and Infernal identified the same set of the 3 rRNAs and 46 tRNAs. Additionally, the ribonuclease P (RNase P), SRP, and tmRNA genes were identified by Infernal. No homolog was found for the non-coding 6S RNA gene. A recent computational screen for 6S RNA across all bacterial phyla (Wehner et al., 2014) reported the absence of 6S RNA in *Pirellula staleyi* and *Rhodopirellula baltica*, which are the closest related species to *T. terrifontis* in the 16S rRNA phylogeny (Slobodkina et al., 2015). This suggests that this gene is also likely to be absent in *T. terrifontis.* The Infernal search also revealed three riboswitches: Cyclic di-GMP-I (RF01051), сobalamin (RF00174), and fluoride (RF01734).

The genome was submitted to GenBank with the accession number CP018477.

### 3.2. Transcriptome sequencing and general transcriptome characteristics

*T. terrifontis* R1 cells were cultured in growth media containing trehalose or xanthan gum, each in triplicates (see Materials and methods). Transcriptome sequencing using Illumina paired-end sequencing resulted in between 11.5m and 12.1m read pairs for the 2 x 3 replicates. Across these, between 91.3% and 98.5% of the read pairs could be mapped uniquely to the genome. Differential expression analysis reported that 665 genes are up- and 617 genes are down-regulated on xanthan gum compared to trehalose-grown culture according to the edgeRun unconditional exact test using a Benjamini-Hochberg corrected P-value < 0.05 (Fig. 1, Table S1).

**Figure 1.**
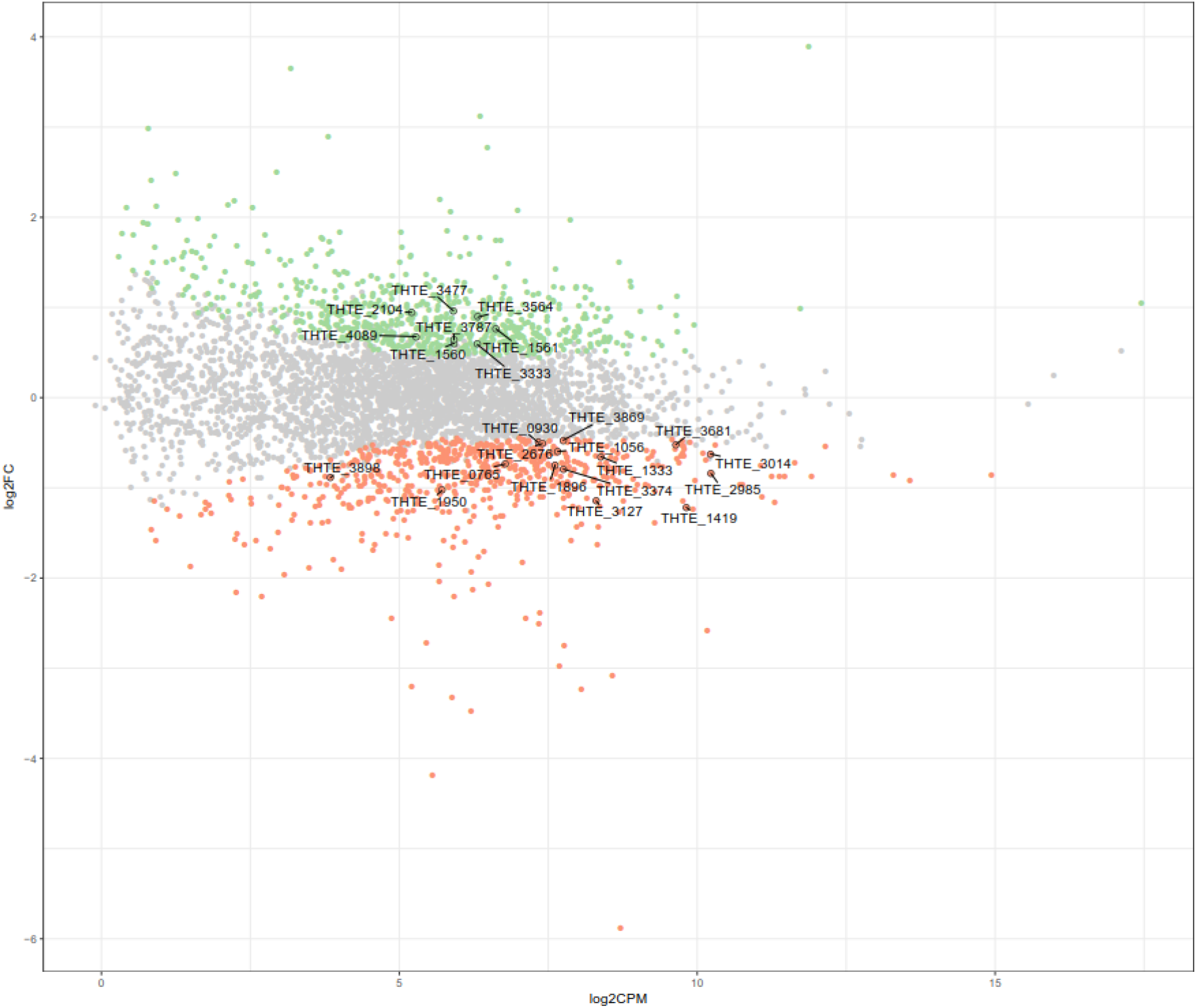
Differently expressed genes between xanthan gum-grown and trehalose-grown cells: green and red dots indicate significantly up-regulated and down-regulated genes in xanthan gum culture compared with trehalose culture (FDR < 0.05).

### 3.3. Genome-scale reconstruction of oligo- and polysaccharide degradation

*T. terrifontis* R1 was shown to be able to grow using the following oligo- and polysaccharides as substrates: sucrose, trehalose, cellobiose, starch, xylan, pectin or xanthan gum. No growth was detected when maltose, lactose, agarose, alginate, cellulose, chitin or inulin were added to the medium as sole carbon sources (Slobodkina et al., 2015).

Our analysis of the *T. terrifontis* R1 genome revealed 101 genes encoding glycosidases (GHs), 14 genes encoding polysaccaride lyases (PLs) and 3 genes encoding carbohydrate esterases (CEs) (Table S2). Among these, 54 genes encode proteins that were predicted to be secreted outside the cells (whether anchored on the cells surface or being released into the culture broth). No dominant CAZy (GH, PL or CE) families were observed among *T. terrifontis* R1 CAZymes, yet the most numerous were GH5 and putative glycosidases, including DUF1080 domain. Detailed analysis of the CAZymes specificities revealed following activities: trehalose can be degraded by trehalose synthase acting in opposite direction (THTE_2039). Sucrose hydrolysis may occur by the action of intracellular fructosidase (THTE_0696). Alpha-1,4-bonds and alpha-1,6-bonds in starch can be hydrolyzed by a number of GH13 and GH77 glycosidases (THTE_1477, THTE_2143, THTE_3153 and THTE_3783), producing maltooligosacchrides and finally D-glucose. Cellobiose can be hydrolyzed by the putative beta-glucosidase (THTE_0963), or one of the GH2 and GH5 glycosidases with currently uncertain function. Xylan can be decomposed to xylooligosaccharides by means of endoxylanases (THTE_2600 and THTE_3961) and to xylose by beta-xylosidases (THTE_0688, THTE_1819, THTE_1884 and THTE_2108). Pectin degradation occurs, most probably, by the action of a pectate lyase (THTE_1993) and several polygalacturonases (THTE_0436, THTE_1516 and THTE_2121), releasing D-galacturonic residues, that are further metabolized to D-glyceraldehyde 3-phosphate and pyruvate (see below). A large number of glycosidases was predicted to be involved in xanthan gum hydrolysis, see section “*Xanthan gum and trehalose utilization pathways, revealed by comparative genomic and transcriptomic analyses*”.

### 3.4. Genome-scale reconstruction of central carbohydrate metabolism

According to the results of genome analysis, the final products of oligo- and polysaccharides decomposition were predicted to comprise glucose, fructose, mannose, xylose, galacturonate and glucuronate. Some of these (glucose, mannose, xylose) as well as galactose were also shown to be used as growth substrates by *T. terrifontis* R1 according to Slobodkina et al., 2015. D-glucose and D-fructose oxidation seems to occur via the Embden-Meyerhof (EM) pathway (Fig. 2, Fig. 3, Table S3). Interestingly, the genome contains four genes encoding phosphofructokinases: one ATP-dependent (THTE_2190) and three pyrophosphate-dependent (THTE_0093, THTE_1056, THTE_2629). Since no genes for fructose-1,6-bisphosphatase were found, and PPi-dependent phosphofructokinases are thought to be reversible, at least one of them should be a part of gluconeogenesis. Additionally, analysis of the nearest characterized homologs supports two of them to be involved in xylose utilization (see below). The Entner-Doudoroff pathway seems to be inoperative due to the absence of the gene encoding 6-phosphogluconate dehydratase, a key enzyme of the pathway. Glucose-1-dehydrogenase, gluconokinase and gluconate dehydratase genes are also absent in the genome.

**Figure 2.**
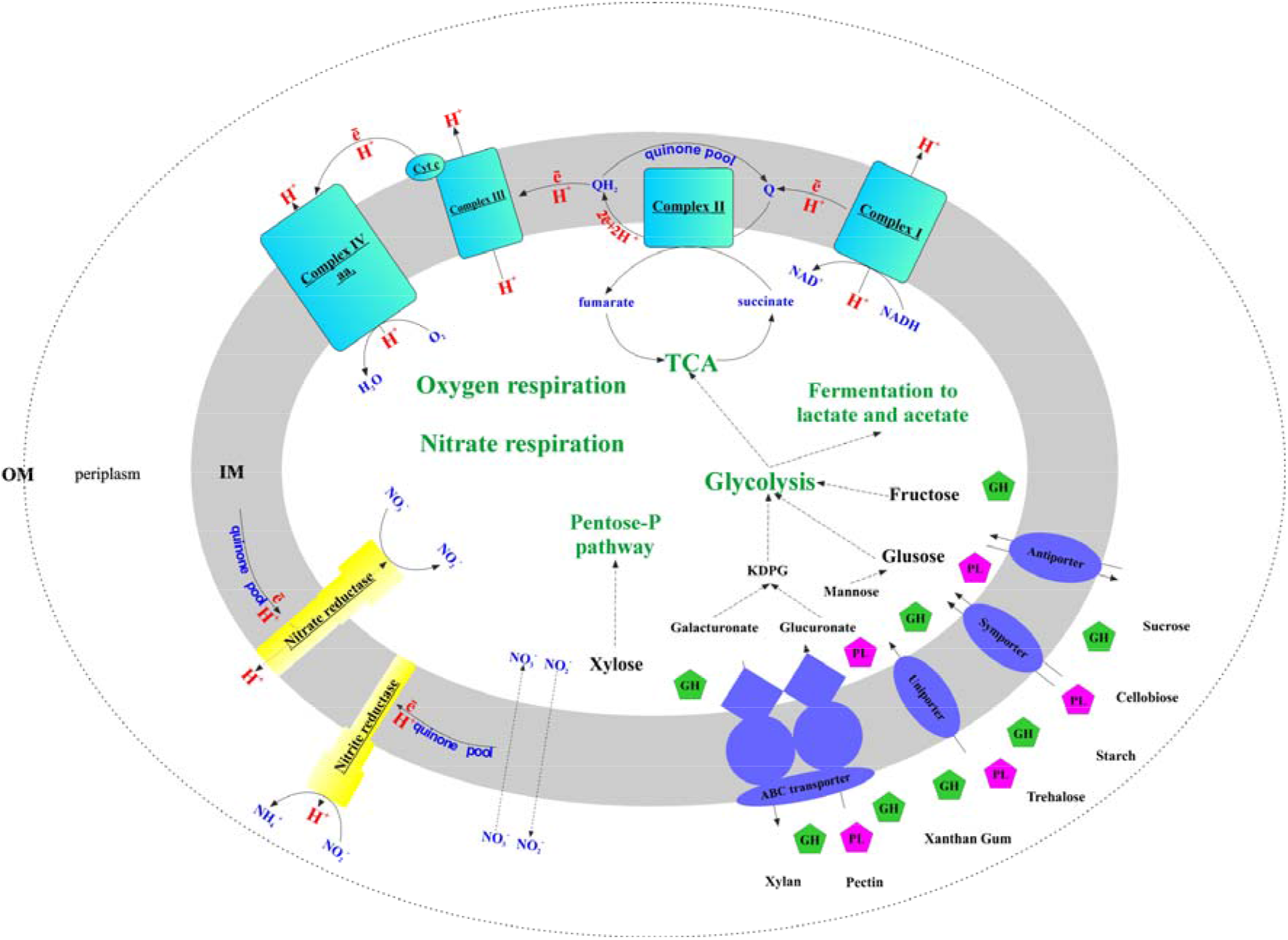
Genomic reconstruction of *T. terrifontis* carbohydrate catabolism, electron transport chain and terminal oxidoreductases: OM, outer membrane; IM, inner membrane; KDPG, 2-keto-3-deoxyphosphogluconate; TCA, tricarboxylic acid cycle; Cyt c, cytochrome *c*; GH, glycosidase; PL, polysaccharide lyase.

**Figure 3.**
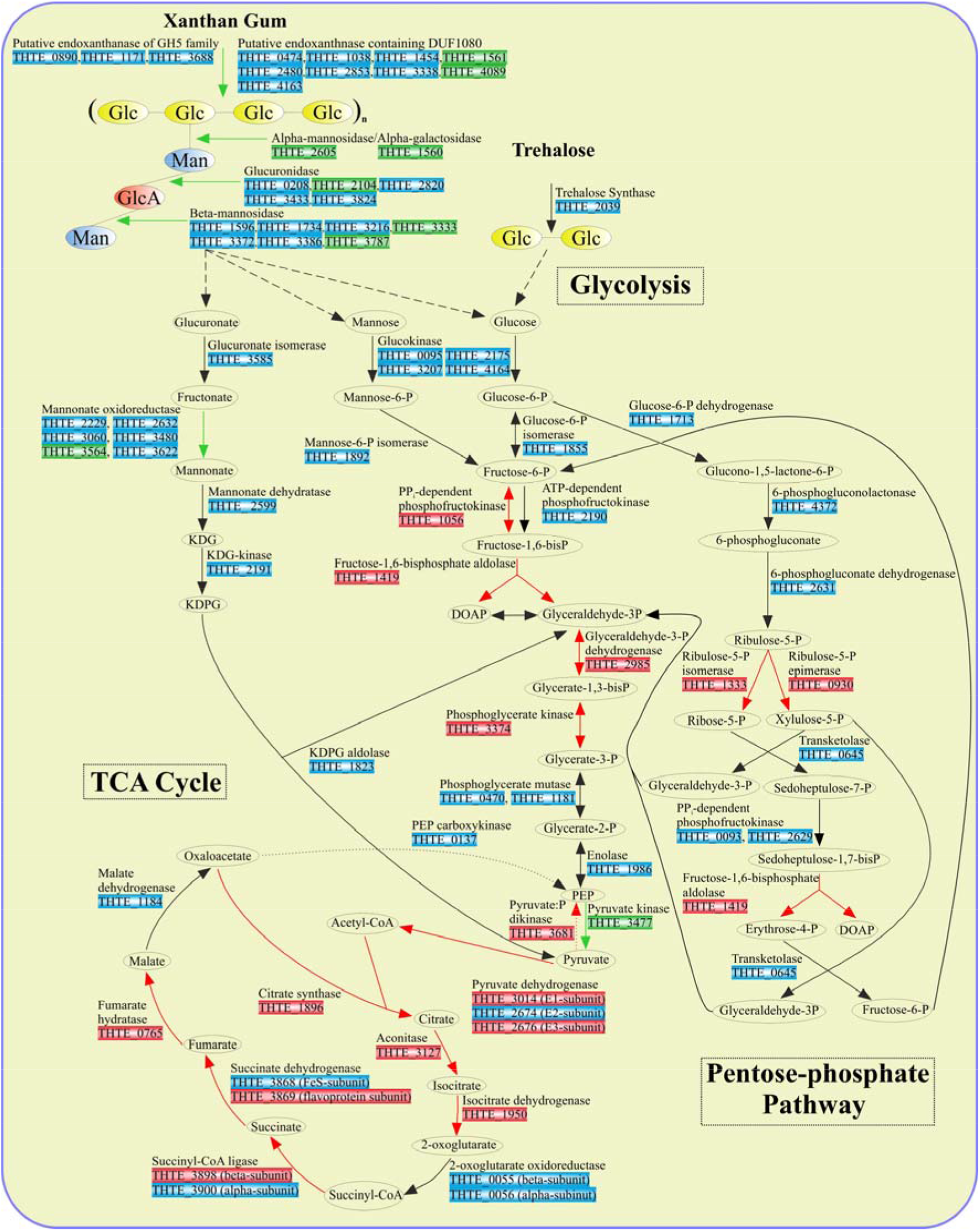
Enzymes involved in sugar turnover. Color boxes indicate the differential expression of the genes, indicated by the transcriptomic analysis. Green: up-regulation with xanthan gum; Red: down-regulation with xanthan gum; Blue: genes are not significantly differentially expressed between xanthan gum and trehalose.

The first step of galactose utilization – phosphorylation to galactose-1-phosphate – is catalyzed by galactokinase (THTE_0177). Next, the putative galactose-1-P-uridyltransferase (THTE_3784) transfers the UDP-group from UDP-glucose to galactose-1-P, producing UDP-galactose and glucose-1-P. This protein belongs to the type 1 galactose-1-P-uridyltransferases family, however, its closest characterized homolog was ADP-glucose:phosphate adenylyltransferase (UniProt ID Q9FK51). While phylogenetic analysis (Fig. S1) supports this finding, the sequence identity of these two proteins is rather low (Identity 35%, Coverage 97%), leaving the function of this enzyme unclear. However, since no other putative galactose-1-P-uridyltransferase genes were found, the assignment of the function to THTE_3784 remains plausible. Finally UDP-galactose is converted to UDP-glucose by the UDP-glucose 4-epimerase (THTE_2863), whereas the glucose-1-phosphate is converted to glucose-6-phosphate by the phosphoglucomutase (THTE_3829).

Xylose utilization was predicted to occur as follows: xylose is isomerized to xylulose by xylose isomerase (THTE_2111); xylulokinase (THTE_0598) phosphorylates xylulose to xylulose-5-phosphate, which finally enters the pentose-phosphate pathway. All genes encoding proteins of both oxidative and synthetic parts of this pathway were found in the genome with transaldolase gene as an exception (Fig. 3, Table S4). Yet, sedoheptulose-7-phosphate (S-7-P), formed under action of transketolase, could be phosphorylated by PPi-dependent phosphofructokinases THTE_0093 and THTE_2629, of which the nearest characterized homolog from *Methylococcus capsulatus* (UniProt Q609I3, Reshetnikov et al., 2008) was shown to reversibly phosphorylate S-7-P with higher activity and affinity than fructose-6-phosphate (F-6-P). The resulting sedoheptulose-1,7-bisphosphate could be eliminated to erythrose-4-phosphate and dihydroxyacetone-phosphate by fructose-1,6-bisphosphate aldolase (THTE_1419) as it was proposed by Susskind et al., 1982 and Schellenberg et al., 2014.

D-galacturonate, released in the course of pectin degradation, is presumably oxidized to glyceraldehyde-3-phosphate and pyruvate through a number of reactions (Fig. S2) catalyzed by uronate isomerase (THTE_3585), putative altronate oxidoreductase (see below), altronate dehydratases (THTE_0455 and THTE_0456), KDG kinase (THTE_2191) and KDPG aldolase (THTE_1823). No genes encoding altronate oxidoreductase belonging to the polyol-specific long-chain dehydrogenase/reductase family (Klimacek et al., 2003) were found. However, the genome contains several genes (*THTE_0865*, *THTE_1784*, *THTE_2229*, *THTE_2632*, *THTE_3060*, *THTE_3480* and *THTE_3564*), probably encoding proteins of the short-chain dehydrogenase/reductase family (J□rnvall et al., 1995). One of its biochemically characterized representatives, an oxidoreductase UxaD from the hyperthermophilic anaerobic bacterium *Thermotoga maritima*, was shown to possess mannonate oxidoreductase activity (Rodionova et al., 2012).

The metabolism of xanthan gum degradation products – glucose, glucuronate and mannose – is described in the section “*Xanthan gum and trehalose utilization pathways, revealed by comparative genomic and transcriptomic analyses*”.

Pyruvate, generated in the course of degradation of sugars and sugar acids, is further oxidized to acetyl-CoA in the reactions catalyzed by the pyruvate dehydrogenase complex (Fig. 3): pyruvate dehydrogenase (E1) (THTE_3014), dihydrolipoamide acetyltransferase (E2) (THTE_2674) and lipoamide dehydrogenase (E3) (THTE_2676).

It has been shown by Slobodkina et al, 2015, that the products of *T. terrifontis* R1 glucose fermentation were hydrogen, lactate and acetate. Lactate could be produced from pyruvate by lactate dehydrogenase (THTE_3348), while the mechanism of acetate formation remains unclear. Although two acetate kinases were found (THTE_1319 and THTE_2274), no genes coding for phosphate-acetyl transferase were detected in the genome. It is therefore possible that acetate could be formed due to the action of CoA-acylating aldehyde dehydrogenase (THTE_1321), catalyzing the NADH-dependent reduction of acetyl-CoA to acetaldehyde (Toth et al., 1999), and aldehyde dehydrogenase (THTE_2212) catalyzing the oxidation of acetaldehyde to acetate along with formation of NADH (Ho and Weiner, 2005). Finally, acetate could be formed under the action of putative ADP-forming acetyl-CoA synthetase (THTE_2996), as it was shown for few hyperthermophilic archaea (Musfeldt et al., 1999; Musfeldt and Schönheit, 2002). Surprisingly, the genome encoded an ATP-dependent acetyl-CoA synthase (THTE_1589), which catalyzes the irreversible activation of acetate, whereas acetate was not listed among the substrates, supporting the growth of *T. terrifontis* R1 in (Slobodkina et al, 2015).

Hydrogen formed by *T. terrifontis* R1 in the course of fermentation apparently results from the operation of group 3c [NiFe]-hydrogenase THTE_4311-4313 and/or [FeFe]-hydrogenases THTE_2884, THTE_2882, THTE_2881, THTE_3842-THTE_3844. The latter enzyme is probably unfunctional or has changed function since the P1 and P2 motifs in its catalytic subunit (Vignais and Billoud, 2007) are strongly impaired and the order of subunit-encoding genes is unusual. On the other hand, according to our analysis, all the genomes of planctomycetes, available in the IMG database (34 genomes of planctomycetes with assigned genus and species names. The analysis was performed 03.07.17), lack genes of [FeFe]-hydrogenase. Since, *T. terrifontis* is the first planctomycete known to synthesize hydrogen in the course of fermentation, and it is currently the only one in which genes for [FeFe]-hydrogenases have been found, at least some of its [FeFe]-hydrogenases could be involved in hydrogen production.

All genes, coding the citrate cycle (TCA cycle) enzymes were found in the *T. terrifontis* R1 genome (Fig. 2, Fig. 3, Table S5).

### 3.5. Genome-scale reconstruction of nitrate reduction

Genes for all three subunits of the respiratory cytoplasmic nitrate reductase Nar (Simon and Klotz, 2013) were found in the genome. The alpha subunit (NarG) THTE_1509 belongs to the deep lineage within the Nar-DMSO cluster (Fig. S3) of the molybdopterine superfamily (Duval et al., 2008). *THTE_1508* and *THTE_1507* encode the other two subunits NarH and NarI, respectively, while *THTE_1506* encodes a chaperon subunit (TorD). The NarGHI complex might form a supercomplex with an electrogenic membrane-bound NADH dehydrogenase (Simon and Klotz, 2013, Fig. 2, Table S6). No diheme subunit NarC was found, yet the genes of b/c1 complex (complex III, *THTE_1510*-*THTE_1512*) are located in close vicinity to the NarGHI genes (*THTE_1509*-*THTE_1507*), what might reflect the involvement of the complex III in the electron and proton transfer during *T. terrifontis* anaerobic growth with nitrate. Nitrite is reduced to ammonium by means of non-electrogenic periplasmic membrane-bound nitrite reductase Nrf, the catalytic subunit NrfA and the membrane-bound subunit NrfH (Simon and Klotz, 2013) of which are encoded by

*THTE_1450* and *THTE_1449*, respectively.

### 3.6. Genome-scale reconstruction of aerobic respiration

The complete aerobic respiratory electron transfer chain (ETC), including H^+^-translocating NADH-dehydrogenase (complex I), succinate dehydrogenase (complex II), cytochrome b/c1-complex (complex III), and terminal cytochrome c oxidase aa3-type (complex IV) was found (Fig. 2, Table S6,). We did not find the typical cytochrome *c* gene or the plastocyanin gene, involved in transferring electrons from complex III to complex IV, yet *THTE_3354* encodes a putative large (258 amino acids) cytochrome *c* containing two monoheme domains. No genes encoding terminal quinol oxidases (bd-type, bo3-type or ba3-type) were found in the genome.

### 3.7. Xanthan gum and trehalose utilization pathways, revealed by comparative genomic and transcriptomic analyses

In order to decipher the mechanisms of xanthan gum degradation, *T. terrifontis* R1 was grown on xanthan gum and trehalose (as the control), and the transcriptomes were sequenced and analyzed for genes that are up-regulated in the cultures with xantham gum as the substrate. Trehalose is a disaccharide consisting of 1-1-alpha-linked glucose molecules, and it was chosen as one of the simple sugars, supporting good growth of the strain. Interestingly, *T. terrifontis* R1 genomic analysis revealed no genes coding for known trehalose-hydrolyzing enzymes of GH15, GH37 and GH65 families. Furthermore two GH13 proteins (THTE_1477 and THTE_3153) have no trehalose-converting enzymes among their nearest characterized relatives. Therefore, the only remaining reasonable candidate involved in decomposition of trehalose is a trehalose synthase of GT4 family (THTE_2039), acting in reverse direction, leading to a release of D-glucose and NDP-D-glucose molecules. The level of its expression in cells, grown on trehalose and xanthan-gum was similar (Fig. 3), what could be explained by reversibility of its action (Qu et al., 2004; Ryu et al., 2005) at various growth conditions: trehalose degradation when trehalose is being sole substrate and trehalose synthesis when other substrates are used.

Despite its ubiquitous usage in pharmaceutical and food industries, not much is known about xanthan gum (beta-1,4-glucan with mannosyl–glucuronyl–mannose side chains) degradation mechanisms. For the complete hydrolysis of the molecule the following linkages should be broken: β-mannose-1–4-α-glucuronate, β-glucuronate-1–2-α-mannose, α-mannose-1–3-β-glucose linkages in side chains, and β-1,4-glucosidic linkages in polyglucose backbone (Fig. 4).

**Figure 4.**
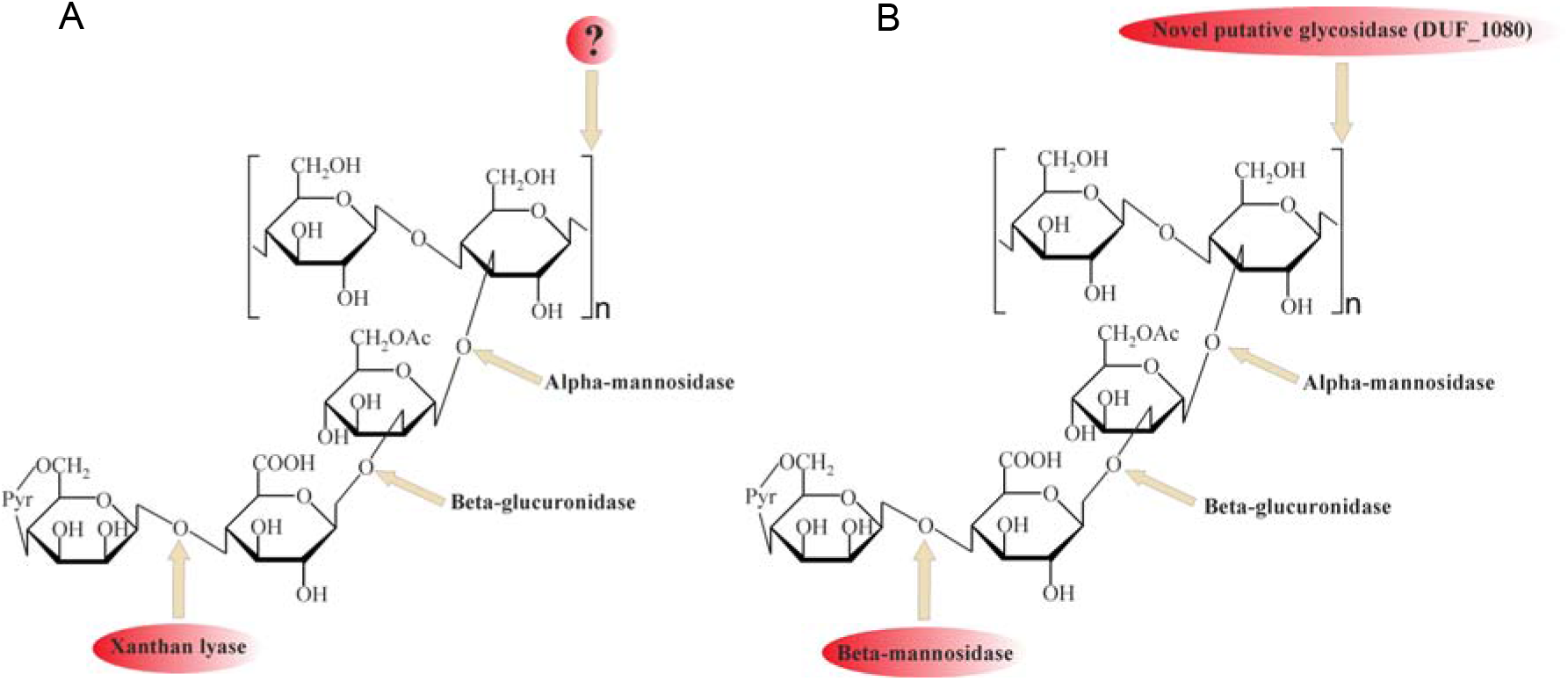
Comparison of (A) traditional (with xanthan lyase) and (B) proposed novel (without xanthan lyase) xanthan gum degradation pathways.

Among the enzymes currently known to be involved in xanthan gum decomposition, there are few removing terminal mannose residues xanthan lyases (Hashimoto et al., 1998; Ruijssenaars et al., 1999), belonging to the PL8 family, and an endoxanthanase (xanthan-specific endoglucanase, Li, Guo et al., 2009), hydrolyzing the glucan backbone(Fig. 4A). The latter has been biochemically characterized, yet its sequence is still unknown, preventing structure analysis and evolutionary reconstructions using sequence comparison.

No homologs of PL8 family lyases, to which all known xanthan lyases belong, were found in the *in silico* translated *T. terrifontis* R1 proteome. They were probably replaced by several putative endomannanases/beta-mannosidases of GH5 family (THTE_1596, THTE_1734, THTE_3216, THTE_3333, THTE_3372, THTE_3386 and THTE_3787, Fig. S4B), cleaving Man(1β–4α)GlcA linkages. Four of them (THTE_1596, THTE_3216, THTE_3333 and THTE_3386) were predicted to be extracellular. Transcriptomic analysis showed two genes *THTE_3333* and *THTE_3787* to be up-regulated in the xanthan gum cultures, assuming their involvement in its decomposition.

For cleavage of the GlcA(β1–2α)Man linkage the action of beta-glucuronidase is needed. All known beta-glucuronidases belong to the GH1, GH2, GH30 and GH79 families. While no genes encoding GH1, GH30 and GH79 proteins were found, five genes encoding GH2 family glycosidases (*THTE_0208*, *THTE_2104*, *THTE_2820*, *THTE_3433* and *THTE_3824*) were revealed in the *T. terrifontis* R1 genome. The family GH2 contains a number of enzymes with various specificities. We compared *T. terrifontis* GH2s with the previously characterized members of the GH2 family, to predict whether the putative five *T. terrifontis* GH2 proteins act as beta-glucuronidases. All five genes formed a monophyletic group, adjoined to the cluster with characterized beta-galactosidases and beta-glucuronidases (Fig. S5). Given that one of these (*THTE_2104*) was significantly up-regulated in the cells growing on xanthan gum, the beta-glucuronidase activity seems to be characteristic of at least this one, yet possibly all five GH2 from this monophyletic group possess this activity. It should be noted that THTE_2104 and also THTE_0208 were predicted to be secreted.

The Man(1α–β3)Glc linkage could be hydrolyzed by an alpha-mannosidase of GH38 family (THTE_2605). Another option is an extracellular putative alpha-galactosidase of GH36 family (THTE_1560), whose characterized homologs are known to hydrolyze a number of oligosaccharides of various structures (Merceron et al., 2012). Both *THTE_2605* and *THTE_1560* genes were up-regulated during the growth on xanthan gum.

Finally, the hydrolysis of a Glc(β1–4)Glc linkage in xanthan gum backbone could be catalyzed by the GH5 enzymes THTE_0890, THTE_1171 and THTE_3688, however, only one of them was predicted to be extracellular (THTE_1171) and none of them were upregulated on xanthan gum.

Search of the other putative glycosidases of *T. terrifontis* R1 revealed nine proteins (THTE_0474, THTE_1038, THTE_1454, THTE_1561, THTE_2480, THTE_2853, THTE_3338, THTE_4089 and THTE_4163) containing a domain of unknown function (DUF1080). All of these proteins, except THTE_1454 and THTE_3338, were predicted to be extracellular, and two of them (*THTE_1561* and *THTE_4089*) were up-regulated on xanthan gum. These proteins may be representatives of a novel family of glycosidases according to TOPSAN annotation (http://www.topsan.org/Proteins/JCSG/4jqt). Such a high number of genes encoding putative glycosidases with unknown function in a xanthan gum degrading microorganism might be an indication on their involvement in the process, most probably for hydrolysis of the backbone linkage (Fig. S4B). Interestingly, proteins containing DUF1080 are highly overrepresented among all *Planctomycetes* (Table S7) in comparison with other organisms including *Bacteroidetes*, where they are also in high number, but in much lower abundance compared with *Planctomycetes.* Finally, five of nine *T. terrifontis* R1 DUF1080 proteins, including THTE_1561, have a CBM66 domain, which was found mainly among *Firmicutes* representatives and helps binding the terminal fructoside residue in fructans (Cuskin et al., 2012). Yet, the DUF1080 and CBM66 domains overlap each other, indicating two different designations of the same domain occurred.

The predicted products of xanthan gum degradation are mannose, glucuronic acid and glucose. Although no mannokinase genes were found in the *T. terrifontis* R1 genome, mannose could be phosphorylated to mannose-6-P by the variety of its putative glucokinases from the ROK family (THTE_0095, THTE_2175, THTE_3207, THTE_4164), representatives of which are known to be capable of acting on various hexoses (Conejo et al., 2010; Nakamura et al., 2012). Mannose-6-phosphate upon conversion to fructose-6-phosphate by mannose-6-phopshate isomerase (THTE_1892) enters the EM pathway (Figs. 2, 3). However, none of these proteins were up-regulated on xanthan gum in our experiment.

An oxidation of glucuronate, released during xanthan gum hydrolysis, might occur through formation of fructonate, followed by reduction to mannonate, dehydration to 2-keto-3-desoxy-6-phosphogluconate (KDPG) and its elimination to pyruvate and glyceraldehyde-3-phosphate (Fig. 3). All genes encoding the respective proteins, except mannonate oxidoreductase, were found in the genome. As it was hypothesized for altronate oxidoreductases (see above), we suggest that some of short-chain reductases/dehydrogenases (THTE_2229, THTE_2632, THTE_3060, THTE_3480, THTE_3564 and THTE_3622) with unknown specificity, especially the up-regulated THTE_3564, may act as a mannonate oxidoreductase.

In most cases, the majority of enzymes involved in both the trehalose and xanthan gum-dependent pathways of the central carbohydrate metabolism were expressed on the same level or were down-regulated in the cells grown on xanthan gum. This could be due to the lower structural complexity of trehalose in comparison with xanthan gum, which requires fewer degradation steps and determines easier import into the cell: one transporter and one enzymatic step are enough to transport and decompose trehalose to the basic metabolites (D-glucose and NTP-α-D-glucose) compared with most certainly multiple transporters and four steps of xanthan gum decomposition, coupled with two- and five-step mannose and glucuronate, respectively, conversions to EM pathway metabolites (Fig. 3). Finally, the majority of flagellar, as well as pili IV and secretion system proteins, were up-regulated (Table S1) in the cells grown on xanthan gum. This might be a reflection of viscosity of the substrate, which force cells to be more agile and capable of binding to the substrate.

## 4. Conclusion

*Thermogutta terrifontis* is the first thermophilic and one of the first anaerobic planctomycetes known so far. Like other planctomycetes, it is capable of growing on various polysaccharides, including xanthan gum, a complex bacterial polysaccharide with many applications in various fields of biotechnology. Our analysis of the complete *T. terrifontis* R1 genome revealed the presence of numerous genes encoding carbohydrate-acting enzymes, including 101 glycosidases genes, 14 polysaccharide lyases genes and 3 carbohydrate esterases genes, which allow this bacterium to use a wide range of sugars, including oligo- and polysaccharides, as sole carbon and energy source. We reconstructed polysaccharide degradation pathways, as well as the pathways of aerobic or anaerobic (nitrate reduction to ammonium) respiration. Complete Embden-Meyerhof-Parnas pathway, TCA cycle, and modified pentose-phosphate pathway (with the absence of transaldolase) were reconstructed. Although our comparative transcriptomics approach has not revealed any peculiarities in the central carbohydrate metabolic traits of cells grown on trehalose or xanthan gum, it helps to establish a novel xanthan gum degradation pathway based on our genome analysis. The proposed pathway involves endomannanases/beta-mannosidases instead of xanthan lyases as well DUF1080 proteins for the hydrolysis of xanthan gum backbone. Surprisingly, the genes coding DUF1080 proteins were highly abundant in *T. terrifontis* R1, as well as in many other *Planctomycetes* genomes. The proposal of the novel pathway of xanthan gum degradation is relevant due to lack of the information on microorganisms degrading xanthan gum and its degradation pathways, which has so far been limited to few representatives of *Actinobacteria* and *Firmicutes* and their enzymes. Yet, further studies including proteomics of xanthan gum-growing cultures, as well as, purification and characterization of the respective enzymes are needed to verify the predicted pathway.

## Data availability

The *T. terrifontis* R1 genome is available via Genbank accession number CP018477.

## Acknowledgements

The work was supported by the European Union 7th Framework Programme FP7/2007-2013 under grant agreement no. 265933 (*Hotzyme*). The work of AE and IK was supported by RSF grant 16-14-00121.

